# MetaResNet: Enhancing Microbiome-Based Disease Classification through Colormap Optimization and Imbalance Handling

**DOI:** 10.64898/2026.03.11.708118

**Authors:** Awais Qureshi, Abdul Wahid, Shams Qazi, Hasan Ali Khattak, Syed Fawad Hussain

## Abstract

Image-based representations of metagenomic data enable convolutional neural network (CNN) applications for microbiome disease classification. However, the impact of colormap selection on model performance remains unexplored. Current approaches arbitrarily select visualization parameters despite evidence that colormap choices can suppress minority-class features in imbalanced microbiome datasets. This study systematically evaluates colormap effects on metagenomic disease classification to establish evidence-based visualization guidelines. We developed MetaResNet, a custom CNN architecture incorporating residual blocks and attention mechanisms, to assess five colormap schemes (Jet, YlGnBu, Reds, Paired and nipy spectral) across four benchmark datasets: inflammatory bowel disease (n=110), colon cancer (n=121), women type 2 diabetes (n=96), and obesity (n=253). Class imbalance was addressed using Synthetic Minority Over-sampling Technique (SMOTE) versus class weighting strategies. Custom data augmentation preserved taxonomic abundance relationships while enhancing generalization. Performance evaluation employed F1-score, Receiver Operating Characteristic and Area Under the Curve (AUC-ROC), Matthews correlation coefficient (MCC), precision, and recall to address accuracy limitations in imbalanced scenarios. Results identified the Jet colormap coupled with SMOTE as the optimal global configuration, maximizing signal retention and achieving peak performance (AUC 1.00 in Colon). SMOTE significantly improved minority-class recall over class weighting (0.81 ± 0.09 vs. 0.69 ± 0.11, *p* = 0.003). MetaResNet achieved performance comparable to current state-of-the-art frameworks, while statistically outperforming established deep learning baselines (e.g., DeepMicro, PopPhy-CNN; *p* = 0.025) in discriminatory power (AUC), with peak values exceeding 0.96. These findings demonstrate that visualization efficacy is strategy-dependent, establishing MetaResNet as a robust framework for microbiome-based diagnostics that supports evidence-based visualization strategies for precision medicine.

## 1 INTRODUCTION

Metagenomics is transforming disease research by enabling comprehensive analysis of the human gut microbiome, an ecosystem harboring 100-fold more genes than the human genome. This genetic diversity provides critical insights into microbial community structure, metabolic pathways, and disease-associated dysbiosis. Metagenomic workflows involve DNA sequencing, taxonomic classification, and functional annotation to convert raw reads into operational taxonomic units (OTUs), which serve as the foundation for microbial diversity analysis and disease classification Liu et al. (2021).

However, data representation strategies—including visualization choices such as colormaps for OTU abundance—remain largely unexplored, despite their potential impact on both model performance and biological interpretability. This issue becomes especially critical in small and imbalanced datasets, where poor color choices can distort abundance gradients, suppress minority-class features, and amplify visual noise.

Recent advances in metagenomic disease classification have increasingly adopted deep learning approaches to address the challenges of high-dimensional, sparse OTU data. Traditional machine learning methods such as Random Forest and Support Vector Machines struggle with the curse of dimensionality inherent in microbiome datasets Oh and Zhang (2020); Volkova and Ruggles (2021). To overcome these limitations, researchers have developed innovative image-based representations that transform one-dimensional OTU abundance vectors into two-dimensional “synthetic images,” enabling the application of CNN Nguyen et al. (2019a, 2020). Notable frameworks include PopPhy-CNN, which leverages phylogenetic structure Reiman et al. (2020), and MAGNAM, which creates multichannel microbiome prints through manifold embedding Shen et al. (2023). While these approaches have demonstrated superior performance over traditional methods, they consistently note the arbitrary selection of visualization parameters—particularly colormap choices—with studies reporting that `viridis` and `Jet` colormaps improve accuracy for specific datasets Nguyen et al. (2019a, 2020).

Despite these advances, critical methodological gaps remain in image-based metagenomic approaches. First, while studies have experimented with different colormaps for OTU visualization Nguyen et al. (2018, 2019a, 2020), no systematic evaluation of colormap effects on model performance been conducted. Second, existing studies exhibit over-reliance on accuracy as the primary performance metric, with insufficient consideration of precision, recall, AUC, and F1-score—particularly problematic given the class imbalance prevalent in microbiome datasets. Third, the impact of colormap selection on minority-class feature representation in imbalanced datasets remains unexplored, despite evidence that visualization choices can suppress critical disease signals Shen et al. (2023).

To address these critical gaps, this study presents the first systematic evaluation of colormap effects on metagenomic disease classification performance. We introduce MetaResNet, a novel CNN architecture integrating residual blocks and attention mechanisms, specifically designed to robustly evaluate colormap impacts across diverse microbiome datasets. Our comprehensive framework addresses four key contributions:

1. Systematic assessment of colormap schemes (including Jet, YlGnBu, Reds, Paired, and nipy spectral) on CNN performance using our proposed MetaResNet architecture with customized attention weighting mechanisms.
2. Detailed analysis of colormap effects on minority-class feature representation in imbalanced microbiome datasets.
3. Comprehensive evaluation using multiple performance metrics (F1-score, precision, recall, AUC-ROC, and MCC) beyond accuracy to provide robust assessment in imbalanced scenarios.
4. Comparative analysis against state-of-the-art DL techniques to demonstrate the significance of systematic colormap selection.

By addressing methodological gaps, our study aims to establish evidence-based guidelines for colormap selection in metagenomic visualization, potentially improving diagnostic accuracy of microbiome-disease relationships in precision medicine applications.

## 2 MATERIAL AND METHODS

### 2.1 Acquisition and preprocessing of metagenomic datasets

We evaluated our methodology on four different datasets, including data estimated at the species level for various diseases: colon cancer (COL) Zeller et al. (2014), obesity (OBE) Le Chatelier et al. (2013), inflammatory bowel disease (IBD) Qin et al. (2010), and WT2D (Women Type 2 Diabetes) Karlsson et al. (2013). A total of 580 human intestinal metagenomic samples were collected from these datasets. The resources generated from raw sequencing data, processed via the MetaPhlAn2 Blanco-Miguez et al. (2022) and HUMAnN2 Beghini et al. (2021) pipelines, are well-documented and easily accessible to the broader scientific community, as noted in Pasolli et al. (2016).

Notably, the distribution of samples across the disease categories is not uniform, resulting in imbalanced datasets. This study employs binary classification to classify the four diseases. A summary of the dataset is provided in Table 1.

**Table 1.** The table provides data from 04 distinct metagenomic datasets. ‘features (1)’ denotes the number of features that represent species abundance. In contrast, ‘features (2)’ signifies the number of features that illustrate the abundance of gene families.

| Datasets | Disease | Disease Samples | Control Samples | Total Samples | Ratio of Patients | Ratio of Controls | features (1) | features (2) |
| --- | --- | --- | --- | --- | --- | --- | --- | --- |
| Colon | *Colon Cancer* | 48 | 73 | 121 | 0.40 | 0.60 | 503 | 1,796,274 |
| IBD | *Inflammatory Bowel Disease* | 25 | 85 | 110 | 0.23 | 0.77 | 443 | 1,730,384 |
| Obesity | *Obesity* | 164 | 89 | 253 | 0.65 | 0.35 | 465 | 1,519,375 |
| WT2D | *Women Type 2 Diabetes* | 53 | 43 | 96 | 0.552 | 0.448 | 381 | 1,415,610 |

### 2.2 Metagenomics Images Generation

This study utilizes the Met2Img methodology Nguyen et al. (2018) to convert (OTU) data into visual images. The methodology transforms OTU abundance values into visual representations through a series of sequential steps. Initially, binning techniques, including Species Bin (SPB) and Quantile Transformation (QTF), categorize abundance values into discrete bins based on their relative abundance. Subsequently, the abundance values are organized into a matrix using the Fill-up technique, which arranges values within a square matrix in a predefined order, such as right-to-left and top-to-bottom. The matrix is visualized using colormaps including `Jet, nipy spectral, Paired, Reds, and YlGnBu`. This process transforms OTU data into synthetic images that represent the abundance of specific microbial taxa. Furthermore, a comprehensive guide for converting OTU abundance tables into images is available at Nguyen et al. (2019b). The converted colormaps of metagenomes, as illustrated in Fig. 1, provide a clear exemplification of this methodology.

**Figure 1.**
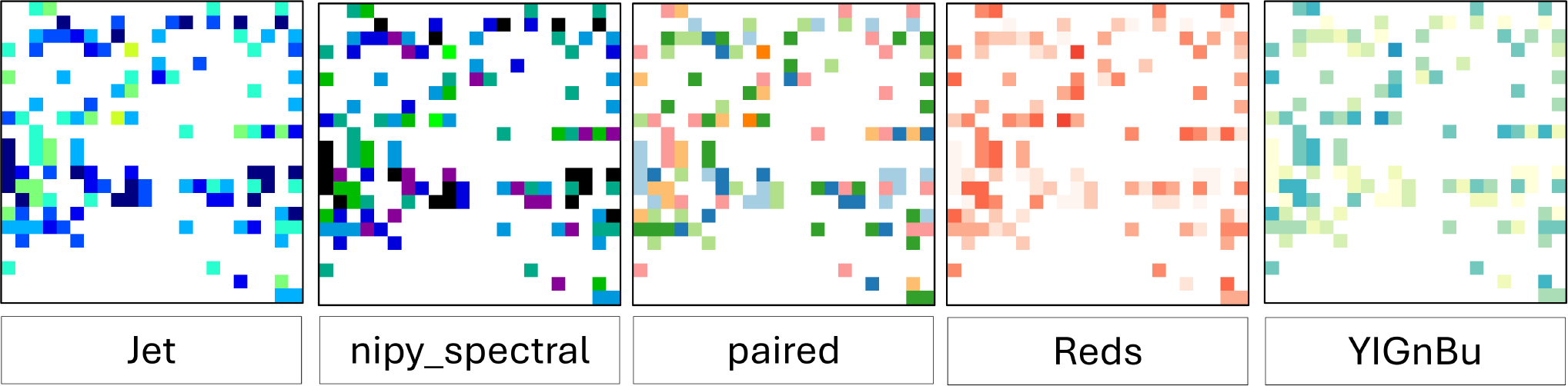
ColorMaps used to represent abundance values in the OTUs, i.e., Jet, nipy spectral, Paired, Reds, and YlGnBu

#### 2.2.1 Colormap Selection Rationale

Foundational studies by Nguyen et al. Nguyen et al. (2018, 2019a, 2020) pioneered the use of synthetic image representations for OTU data, benchmarking standard colormaps to optimize classification performance. Their findings established distinct paradigms based on data distribution: perceptually uniform maps like *Viridis* were optimal for uniform distributions (Quantile Transformation), while spectral maps like *Jet* and *Rainbow* favored bell-shaped distributions (Species Binning). Notably, these studies characterized discrete schemes like *Paired* and *nipy spectral* as suboptimal due to their lack of perceptual uniformity, which was posited to introduce misleading visual artifacts.

This study explicitly challenges that consensus in the context of deep learning. We hypothesized that for sparse, highly imbalanced microbiome datasets, the attribute previously considered a flaw—perceptual non-uniformity—serves as a functional advantage for Convolutional Neural Networks (CNNs). By introducing distinct visual boundaries rather than smooth gradients, discrete schemes like *Paired* potentially amplify the feature distinctness of minority-class OTUs that would otherwise be blended into the background by uniform maps like *Viridis*. Thus, our selection deliberately includes these ‘suboptimal’ schemes to determine if deprioritizing perceptual uniformity enhances CNN sensitivity in imbalanced disease classification scenarios.

### 2.3 Proposed MetaResNet Framework

#### 2.3.1 Initial Model

This study presents the application of a CNN for disease classification from images, as shown in Fig. 2. Prior to the implementation of data augmentation techniques, the research prioritized leveraging the CNN’s structured layer sequence to maximize feature extraction, a critical step for differentiating disease states.

**Figure 2.**
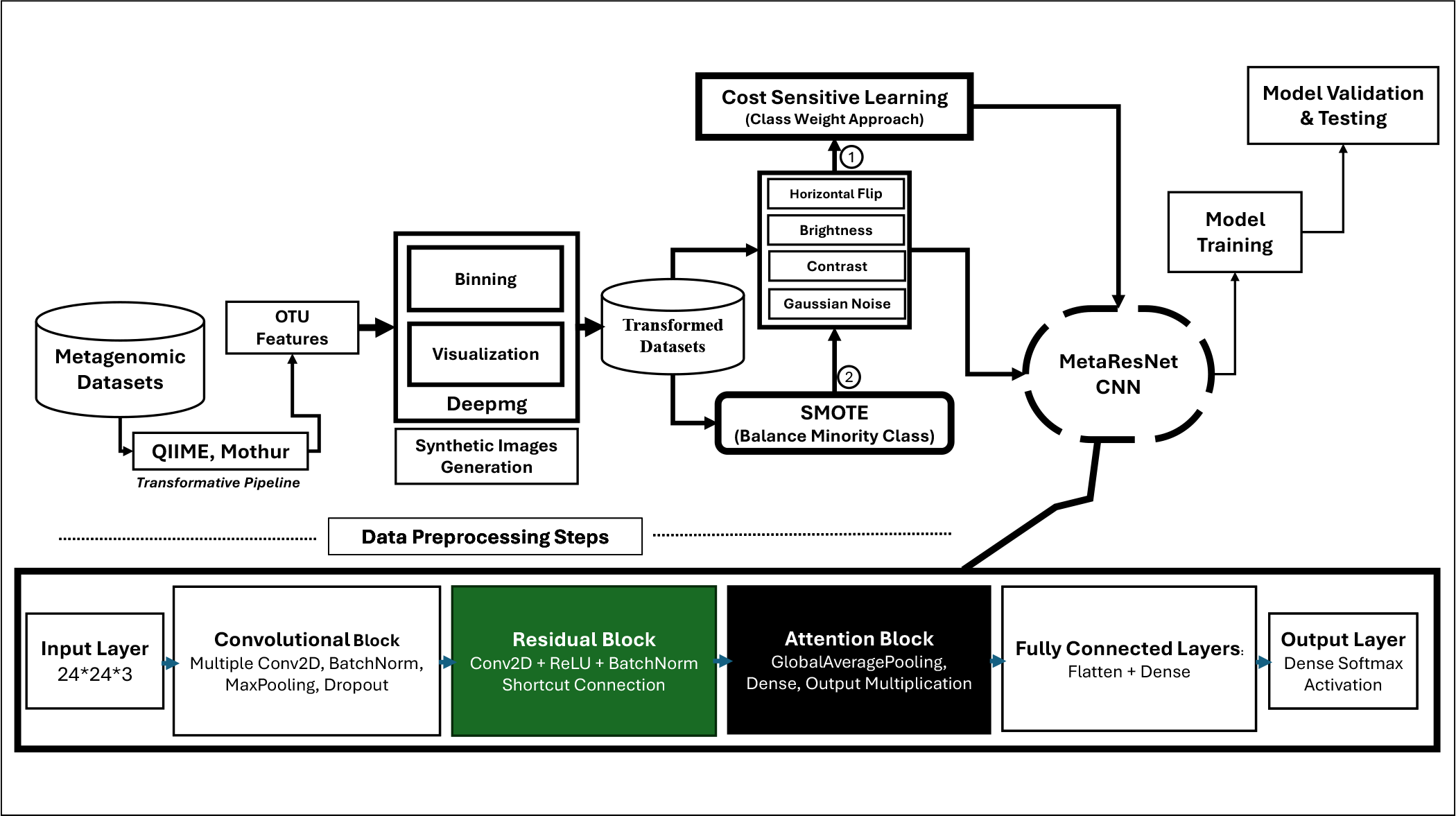
MetaResNet Framework: The MetaResNet Framework architecture outlines the workflow for classifying metagenomic data. The process begins with raw metagenomic datasets, which undergo preprocessing and model training. Input data is transformed using Quantitative Insights Into Microbial Ecology (QIIME) and Mothur to generate (OTU) features. Visualization and synthetic image generation are performed using the Deepmg framework. To address class imbalance, two methods are implemented: cost-sensitive learning using class weights during model training and the Synthetic Minority Over-sampling Technique (SMOTE) prior to training to create balanced datasets. The processed data is then analyzed by the MetaResNet CNN model, which comprises a Convolutional Block, Residual Block, Attention Block, and Fully Connected Layers. Each component is structured to improve feature extraction and classification accuracy, culminating in model validation and testing.

The CNN architecture is illustrated in Algorithm 1. Model training and validation were performed using the computational resources available in the Google Colab environment. Training was executed over 200 epochs with a batch size of 8 and input images of 24 × 24 pixels. The implemented sequential model comprised three convolutional layers with max pooling, followed by a flattening operation, a dense layer with 128 units and Rectified Linear Unit (ReLU) activation, and a final dense layer with Softmax activation for multi-class classification. The Adaptive Moment Estimation (Adam) optimizer was applied with a learning rate of 0.001. Model performance was monitored on the validation set throughout the training process.

Evaluation metrics as defined in the subsection 2.6 included the F1-score, confusion matrix, classification report, and area under the receiver operating characteristic curve (AUC-ROC) to assess class discrimination. This structured approach highlights the architecture’s feature extraction capabilities and introduces measures

##### Algorithm 1

Pre-Augmentation CNN Training Process

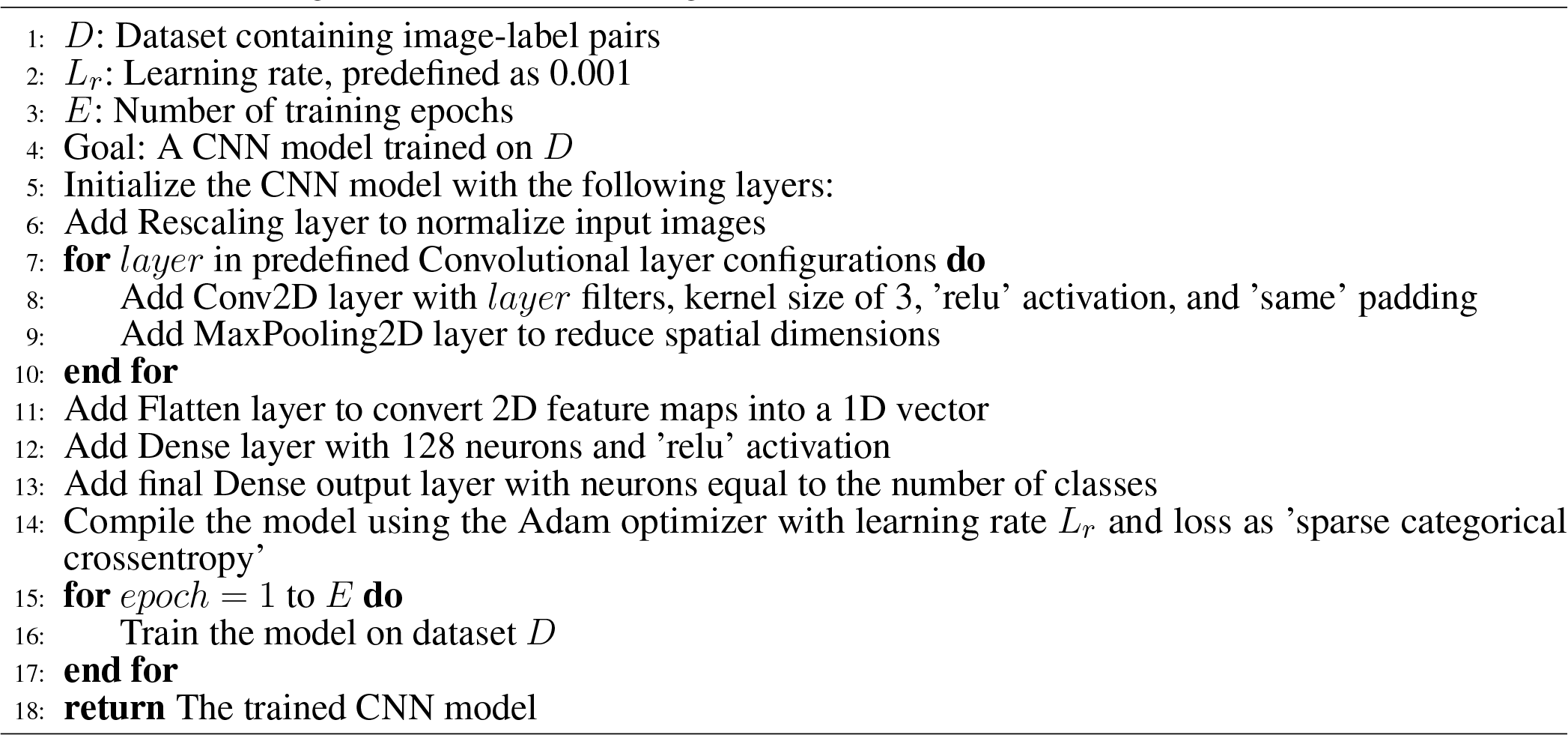

#### 2.3.2 Augmented Model

Data augmentation was applied to increase dataset diversity and improve model robustness. The protocol included random horizontal flipping, which preserves the spatial orientation of abundance patterns, as well as slight modifications in brightness and minor adjustments to contrast to introduce variability without altering image structure or relative abundance. Small amounts of Gaussian noise were added to further enhance variability and support model generalization while maintaining key features. The augmented dataset was then used for subsequent model development. A modified model architecture was developed using the augmented dataset, as shown in Algorithm 2. This architecture incorporates residual blocks with 64 filters and an attention mechanism to improve feature representation and learning. The model begins with a convolutional layer containing 32 filters, followed by a residual block, an attention block, and a dense layer with 256 units before the output layer. Training was performed using the Adam optimizer with a learning rate of 0.001 and sparse categorical cross-entropy loss. Early stopping and model checkpointing were implemented to mitigate overfitting.

##### Algorithm 2

MetaResNet with Attention and Data Augmentation

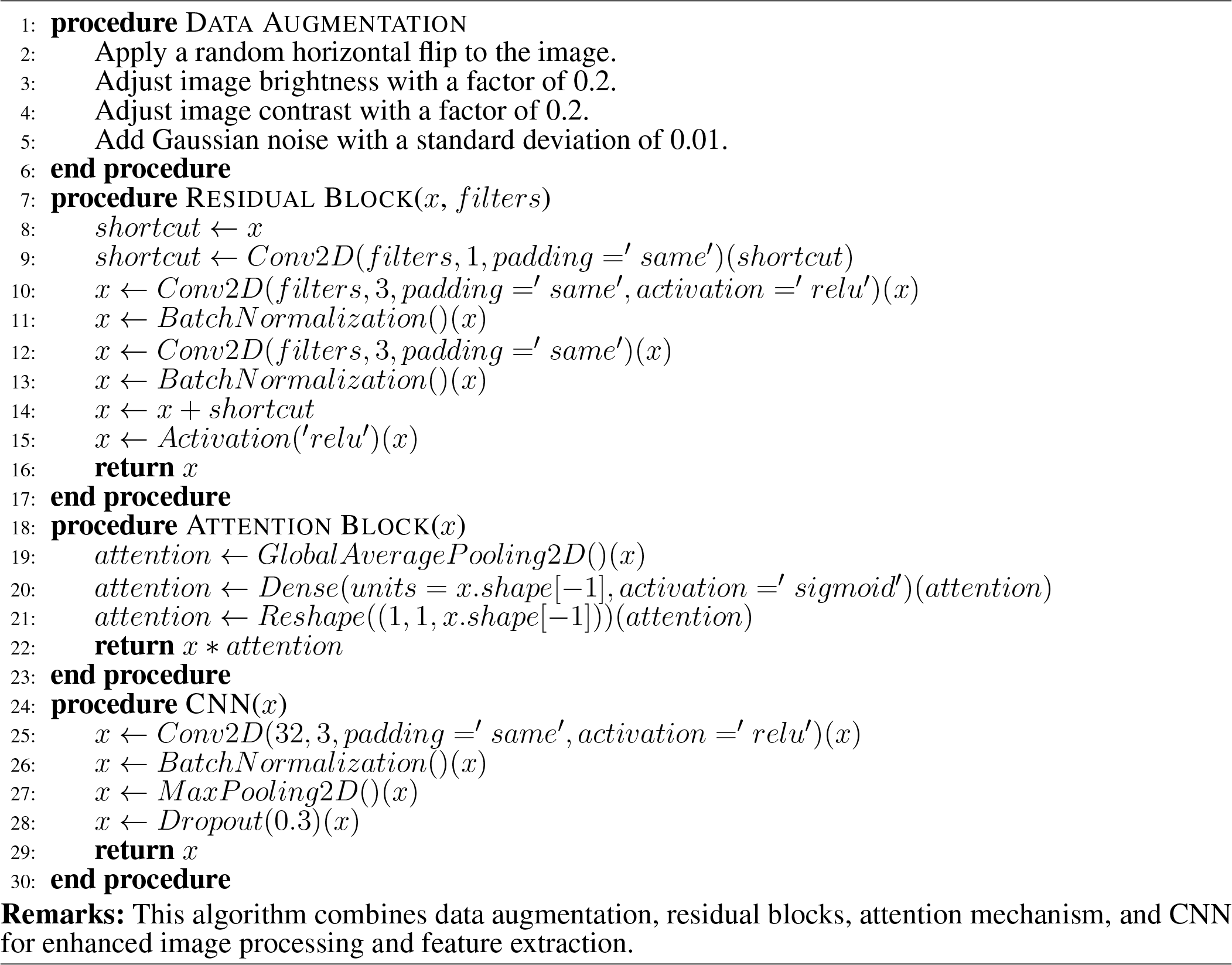

### 2.4 Class Imbalance Handling

To address the critical limitation of class imbalance, which existing deep learning frameworks such as DeepMicro, EnsDeepDP, and PopPhy-CNN generally mitigate only indirectly, we adopted a dual strategy combining SMOTE for synthetic minority oversampling and Class Weights for loss rebalancing. This approach ensures that the minority disease classes are adequately represented during training while simultaneously penalizing their misclassification more heavily, thereby reducing bias toward the majority control class. By explicitly integrating these imbalance-handling techniques into the training pipeline, our framework directly addresses a well-recognized research gap, enhancing both the stability and generalizability of predictions across diverse microbiome datasets.

Both techniques are designed to enhance the model’s generalization capabilities across all classes. The class weights approach stabilizes the training process by ensuring that gradient updates reflect the importance of minority classes. The adapted SMOTE technique provides additional, carefully generated training examples for underrepresented classes, which may enable the model to learn more robust features for these classes without introducing artifacts or distortions that could negatively impact learning.

#### 2.4.1 Cost-sensitive Learning via Class Weights

Class weights as shown in the Equation 1 were computed using the compute class weight function from the sklearn.utils.class weight module King and Zeng (2001). This function calculates weights that are inversely proportional to the class frequencies in the dataset. For one-hot encoded labels, labels were converted to integer class values before calculating the weights. The resulting class weights were formatted as a dictionary, mapping each class index to its corresponding weight. During training, these weights were applied to the loss function via the class weight parameter in the model.fit method. This approach ensures equitable attention to underrepresented classes, resulting in a balanced influence of each class on the loss function.

Class weights (*w*_*j*_) are derived based on class frequencies to address class imbalance, calculated as:

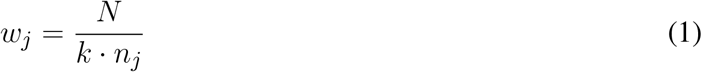

with *N* being the total number of samples, *k* the number of unique classes, and *n*_*j*_ the number of samples for each class *j*. We used the class weights method to handle class imbalance because of its ease of implementation, compatibility with metagenomic classifiers, and interpretability, making it ideal for our framework.

#### 2.4.2 Synthetic Minority Over-sampling Technique (SMOTE) for Images

To address class imbalance, the Synthetic Minority Over-sampling Technique (SMOTE) Chawla et al. (2002) was applied. Unlike standard computer vision tasks where linear interpolation can degrade spatial semantics (e.g., blurring edges in natural photographs), its application here is grounded in the specific nature of omics data. We implemented a three-step process: First, the 2D synthetic images were transformed into 1D feature vectors, effectively treating the pixel grid as a high-dimensional tabular dataset. Second, SMOTE was applied to these *flattened feature representations* to generate synthetic samples for the minority class. Finally, the synthetic vectors were reshaped back to their original image dimensions.

##### 2.4.2.1 Rationale of SMOTE on Synthetic Images

Crucially, because our inputs are visual encodings of biological abundance rather than natural photographs, the linear interpolation performed by SMOTE is mathematically equivalent to averaging the microbial profiles of phenotypically similar patients. This aligns with validated biological augmentation strategies such as *MixUp* and *PhyloMix* Park et al. (2025), which demonstrate that interpolating microbiome vectors preserves community-level patterns. Furthermore, recent studies have successfully employed SMOTE in the feature space for high-dimensional microbiome classification Hong et al. (2025), confirming that this ‘active’ data generation yields robust decision boundaries without introducing the semantic artifacts associated with natural image interpolation.

##### 2.4.2.2 Algorithm Details

The SMOTE process operates as follows:

1. **Identify Nearest Neighbors:** For each minority class sample *x*_*i*_, identify its *k* nearest neighbors from the minority class.
2. **Generate Synthetic Samples:** Randomly select one neighbor *x*_*zi*_ and generate a new sample *x*_*new*_ using Equation 2:

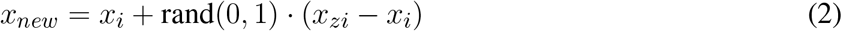

where rand(0, 1) represents a random number between 0 and 1.

Experiments were conducted to rigorously compare this *active* augmentation strategy against *passive* cost-sensitive learning (Class Weights). While Class Weights adjust the loss function to penalize minority misclassifications, SMOTE directly modifies the data distribution by populating sparse regions of the feature space with biologically plausible synthetic samples. Our results as shown in the SubSection 3.2 evaluate whether this feature-space expansion offers superior sensitivity for disease detection compared to loss-weighting alone.

### 2.5 Experimental Setup and Reproducibility

To ensure the rigorous reproducibility of our results and enable valid comparisons between visualization schemes, we implemented a Two-Phase Validation Strategy. This section outlines Phase 1: Model Selection (internal ablation). The protocol for Phase 2: Benchmarking against SOTA methods is detailed in Section 3.5.

#### 2.5.0.1 Data Partitioning

The dataset was partitioned using a **Hold-Out Validation** strategy with a fixed ratio of **57% Training, 27% Validation, and 16% Testing**. This specific partitioning was enforced using a fixed shuffling seed (‘seed=123’) within the TensorFlow `image_dataset_from_directory` pipeline.

- **Training Set:** Used for gradient updates and weight optimization.
- **Validation Set:** Used for hyperparameter tuning and model checkpointing (saving the best weights based on validation loss).
- **Test Set:** A completely unseen hold-out set used strictly for the final performance metrics (MCC, F1-Score) reported in the results.

#### 2.5.0.2 Reproducibility Protocols

All experiments were conducted with fixed random seeds for the operating system, NumPy, and TensorFlow backends (‘random state=42’). This ensured that all models processed the exact same sample indices, ensuring that observed performance differences were attributable to the architectural changes rather than random variations in data splitting.

#### 2.5.0.3 Statistical Analysis

The statistical significance of the observed performance improvements was evaluated using a paired two-tailed Student’s t-test at a 5% significance level (*α* = 0.05). This parametric test was selected because the comparison involved models evaluated on identical cross-validation folds, creating a paired dependency between performance observations. The analysis tested the following hypothesis:

- **Null Hypothesis (***H*_0_): There is no significant difference in the mean performance metrics between the two methods (mean difference = 0).
- **Alternative Hypothesis (***H*_1_): There is a significant difference in the mean performance metrics (mean difference ≠ 0).

The degrees of freedom (*df* ) were calculated based on the number of cross-validation folds (*n* − 1). We assumed that the differences between the paired performance metrics follow an approximately normal distribution, satisfying the assumptions required for the paired t-test.

### 2.6 Evaluation Metrics

The performance of our CNN model is quantitatively assessed using several standard evaluation metrics, such as:

#### Accuracy

Accuracy represents the proportion of true results (both true positives and true negatives) in the total number of cases examined. It is defined as:

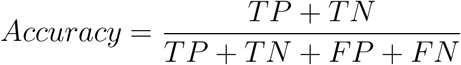

Where *TP, TN, FP*, and *FN* denote the numbers of true positives, true negatives, false positives, and false negatives, respectively.

#### 2.6.1 Precision

Precision, or the positive predictive value, measures the accuracy of positive classifications, calculated as:

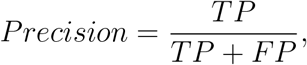

This indicates the model’s ability to identify only relevant instances as positive.

#### 2.6.2 Recall

Recall (or Sensitivity) quantifies the model’s capacity to identify all actual positives, computed as: correctly

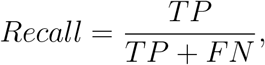

Highlighting its effectiveness in capturing positive instances.

### 2.6.3 F1-Score

The F1-score harmonizes Precision and Recall into a single metric, offering a balanced measure of a model’s precision and recall capabilities, particularly valuable in imbalanced datasets:

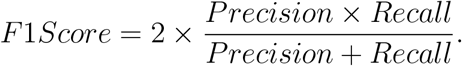

#### 2.6.4 ROC(AUC)

The ROC curve provides a visual representation of a model’s diagnostic ability by plotting the True Positive Rate (TPR) against the False Positive Rate (FPR) across various threshold settings. The AUC quantifies the overall performance, offering a single scalar value that reflects the model’s ability to distinguish between positive and negative instances.

#### 2.6.5 MCC

The MCC considers true and false positives and negatives, providing a balanced measure even in cases of imbalanced class distributions. The MCC is defined mathematically as:

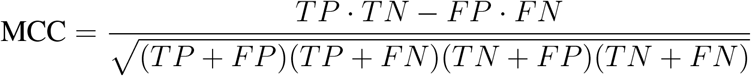

where *TP* represents True Positives, *TN* represents True Negatives, *FP* represents False Positives, and *FN* represents False Negatives. The MCC value ranges from -1 to +1, with +1 indicating perfect prediction, 0 indicating random prediction, and -1 indicating total disagreement between prediction and observation.

## 3 RESULTS

The results of the CNN approach to image-based microbiome disease classification are presented herein, addressing the previously posed research questions. Our findings directly inform the hypotheses regarding the efficacy of CNNs in microbiome data analysis and their potential for enhanced performance and generalization compared to traditional methodologies.

Following the description of the methodological approach, the quantitative results of the CNN model for microbiome disease classification are presented. Table 2 displays the performance metrics of CNN models utilizing the class weighting strategy and Synthetic Minority Over-sampling Technique (SMOTE) across four datasets and five color schemes.

**Table 2.** Comparison of SMOTE and Class Weights Approach on 4 Datasets and 5 Color Schemes.

| Color | Data | Class | SMOTE Approach |  |  |  |  | Class Weights Approach |  |  |  |  |
| --- | --- | --- | --- | --- | --- | --- | --- | --- | --- | --- | --- | --- |
|  |  |  | Prec | Rec | Acc | F1-M | F1-W | Prec | Rec | Acc | F1-M | F1-W |
| Jet | IBD | IBD<br>Ctrl | 0.75<br>0.93 | 0.60<br>0.96 | 0.91 | 0.81 | 0.90 | **1.00**<br>**0.93** | **0.71**<br>**1.00** | **0.94** | **0.90** | **0.93** |
|  | Colon | Col<br>Ctrl | **1.00**<br>**1.00** | **1.00**<br>**1.00** | **1.00** | **1.00** | **1.00** | 0.92<br>0.89 | 0.86<br>0.94 | 0.91 | 0.90 | 0.91 |
|  | WT2D | WT2D<br>Ctrl | **0.92**<br>**1.00** | **1.00**<br>**0.92** | **0.96** | **0.96** | **0.96** | 0.80<br>0.67 | 0.80<br>0.67 | 0.75 | 0.73 | 0.75 |
|  | Obe | Obe<br>Ctrl | 0.75<br>0.75 | **0.89**<br>0.53 | **0.86** | **0.82** | **0.86** | **0.82**<br>0.75 | 0.88<br>**0.65** | 0.80 | 0.77 | 0.79 |
| YlGnBu | IBD | IBD<br>Ctrl | 1.00<br>0.93 | 0.67<br>1.00 | 0.94 | 0.88 | 0.93 | **1.00**<br>**0.96** | **0.80**<br>**1.00** | **0.97** | **0.94** | **0.97** |
|  | Colon | Col<br>Ctrl | 0.75<br>**0.95** | **0.90**<br>0.86 | 0.88 | 0.86 | 0.88 | **0.92**<br>0.89 | 0.86<br>**0.94** | **0.91** | **0.90** | **0.91** |
|  | WT2D | WT2D<br>Ctrl | **0.90**<br>**1.00** | 0.82<br>**0.92** | **0.88** | **0.87** | **0.87** | 0.74<br>0.80 | **0.93**<br>0.44 | 0.75 | 0.70 | 0.73 |
|  | Obe | Obe<br>Ctrl | **0.83**<br>**0.75** | **0.93**<br>0.50 | **0.81** | **0.74** | **0.80** | 0.77<br>0.56 | 0.73<br>**0.61** | 0.69 | 0.67 | 0.69 |
| Red | IBD | IBD<br>Ctrl | **1.00**<br>**0.96** | **0.83**<br>**1.00** | **0.97** | **0.95** | **0.97** | 0.67<br>**0.96** | 0.80<br>0.93 | 0.91 | 0.84 | 0.91 |
|  | Colon | Col<br>Ctrl | 0.89<br>0.91 | 0.80<br>0.95 | 0.91 | 0.89 | 0.90 | **0.93**<br>**0.94** | **0.93**<br>**0.94** | **0.94** | **0.94** | **0.94** |
|  | WT2D | WT2D<br>Ctrl | **1.00**<br>0.76 | 0.64<br>**1.00** | 0.88 | 0.87 | 0.88 | **0.90**<br>**0.93** | **0.90**<br>**0.93** | **0.92** | **0.91** | **0.92** |
|  | Obe | Obe<br>Ctrl | **0.83**<br>**0.82** | **0.91**<br>0.50 | **0.83** | 0.75 | **0.81** | 0.81<br>0.71 | 0.85<br>**0.65** | 0.78 | **0.76** | 0.78 |
| Paired | IBD | IBD | **1.00** | **0.88** | **0.97** | **0.96** | **0.97** | 0.75 | 0.86 | 0.91 | 0.87 | 0.91 |

Table 2 – continued from previous page
| Color | Data | Class | SMOTE Approach |  |  |  |  | Class Weights Approach |  |  |  |  |
| --- | --- | --- | --- | --- | --- | --- | --- | --- | --- | --- | --- | --- |
|  |  |  | Prec | Rec | Acc | F1-M | F1-W | Prec | Rec | Acc | F1-M | F1-W |
|  |  | Ctrl | 0.96 | **1.00** |  |  |  | 0.96 | 0.92 |  |  |  |
|  | Colon | Col<br>Ctrl | **1.00**<br>0.96 | 0.90<br>**1.00** | **0.97** | **0.96** | **0.97** | 0.86<br>**1.00** | **1.00**<br>0.90 | 0.94 | 0.94 | 0.94 |
|  | WT2D | WT2D<br>Ctrl | **1.00**<br>0.76 | 0.64<br>**1.00** | 0.83 | 0.82 | 0.83 | 0.83<br>**1.00** | **1.00**<br>0.86 | **0.92** | **0.92** | **0.92** |
|  | Obe | Obe<br>Ctrl | **0.85**<br>0.69 | **0.89**<br>0.61 | **0.81** | 0.76 | **0.81** | 0.82<br>**0.75** | 0.88<br>**0.65** | 0.80 | **0.77** | 0.79 |
| nipy_spectral | IBD | IBD<br>Ctrl | **1.00**<br>**0.96** | **0.88**<br>**1.00** | **0.97** | **0.96** | **0.97** | 1.00<br>0.89 | 0.57<br>1.00 | 0.91 | 0.84 | 0.90 |
|  | Colon | Col<br>Ctrl | 0.84<br>0.94 | **0.93**<br>0.89 | 0.91 | 0.91 | 0.91 | **1.00**<br>0.95 | 0.92<br>**1.00** | **0.97** | **0.97** | **0.97** |
|  | WT2D | WT2D<br>Ctrl | **1.00**<br>0.81 | 0.73<br>**1.00** | 0.83 | 0.83 | 0.83 | 0.91<br>**1.00** | **1.00**<br>0.93 | **0.96** | **0.96** | **0.96** |
|  | Obe | Obe<br>Ctrl | 0.87<br>0.71 | **0.89**<br>0.67 | 0.83 | 0.78 | 0.83 | **0.88**<br>**0.78** | 0.88<br>**0.78** | **0.84** | **0.83** | **0.84** |

### 3.1 Impact of Colormap Selection on Classification Accuracy

In a comparative analysis of color schemes for metagenomic data visualization, the `nipy_spectral` scheme demonstrated the most consistent stability across the four datasets examined (IBD, Colon, WT2D, and Obesity). This scheme yielded robust accuracy scores, ranging from 83% to 97% for the SMOTE approach and 84% to 97% for the Class Weights approach. Notably, `nipy_spectral` achieved a peak accuracy of 97% for the Colon dataset using Class Weights, while maintaining a strong 91% accuracy with the SMOTE strategy.

The `Red` scheme emerged as the second most effective, exhibiting stable performance with accuracy scores ranging from 83% to 97% for SMOTE and 78% to 94% for Class Weights. Both `nipy_spectral` and `Red` schemes consistently maintained F1 macro and weighted average scores above 80%, indicating balanced feature representation across classes.

Other color schemes showed variable efficacy depending on the balancing strategy. The `Paired` scheme excelled in IBD and Colon datasets, achieving 97% accuracy with SMOTE in both, but demonstrated lower performance in Obesity 81%. Similarly, the `Jet` scheme showed dataset-specific strengths; it achieved perfect classification 100% accuracy for the Colon dataset and superior performance on WT2D 96% using SMOTE. However, its performance dropped significantly when using Class Weights (e.g., 75% on WT2D), highlighting a dependency on synthetic oversampling to resolve decision boundaries.

These findings suggest that while `Jet` may offer peak performance for specific SMOTE-augmented tasks, `nipy_spectral` provides the most reliable generalizability across different imbalance-handling strategies. See the Table 2 for more details.

### 3.2 Efficacy of Imbalance Handling Strategies (SMOTE vs. Class Weights)

To further evaluate the model’s discriminative capability, we analyzed Receiver Operating Characteristic (ROC) curves. Figures 3 and 4 present the class-specific trajectories for the Class Weights and SMOTE strategies, respectively. These plots confirm the model’s robustness, demonstrating high sensitivity and specificity across varying decision thresholds.

**Figure 3.**
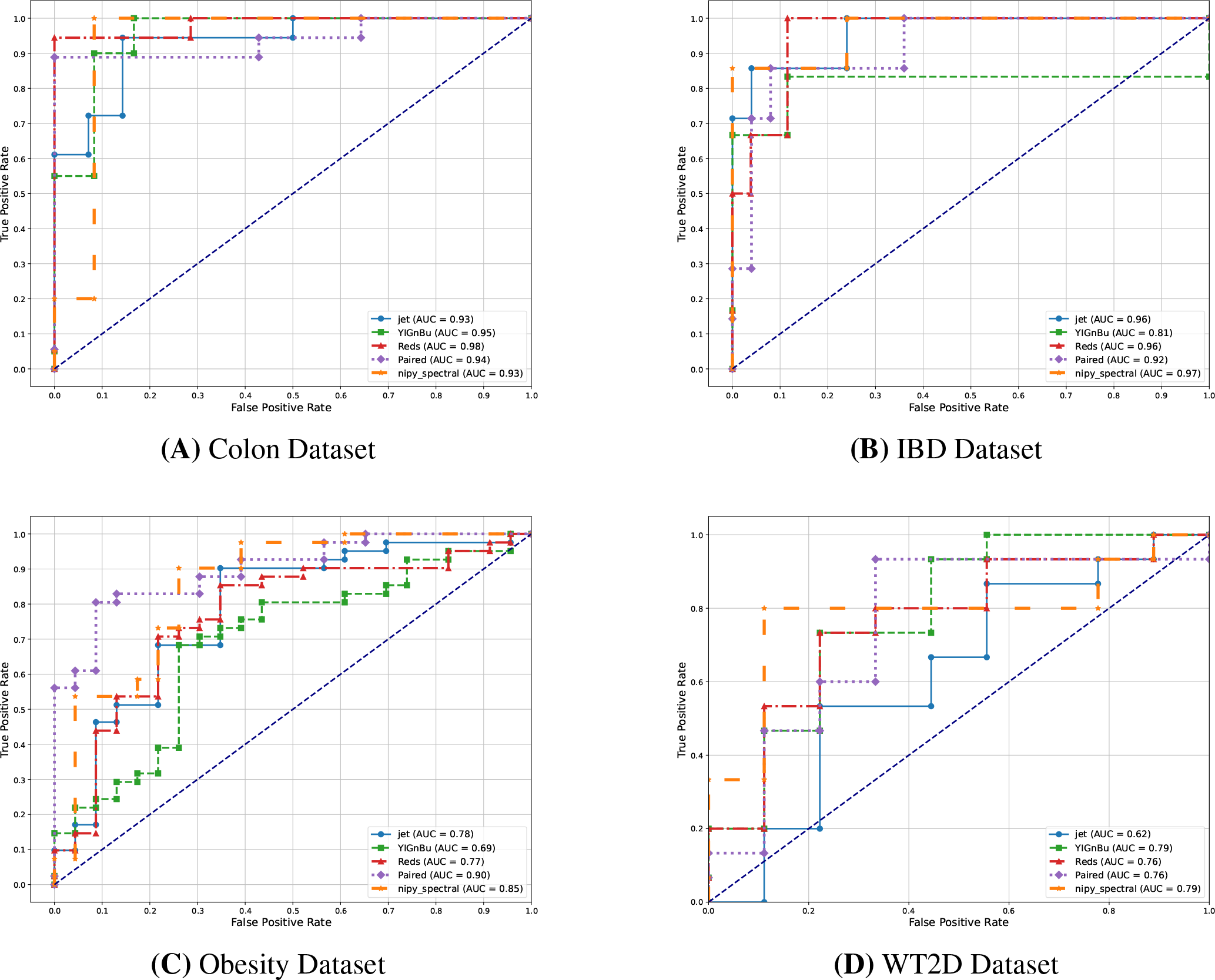
ROC curves for IBD, Colon, wt2d, and Obesity datasets using the Class Weights approach across five color schemes. The figure highlights strong performance in the Colon and IBD datasets, while the wt2d and Obesity datasets show greater variability in AUC values. (A) Colon Dataset. (B) IBD Dataset. (C) Obesity Dataset. (D) WT2D Dataset.

**Figure 4.**
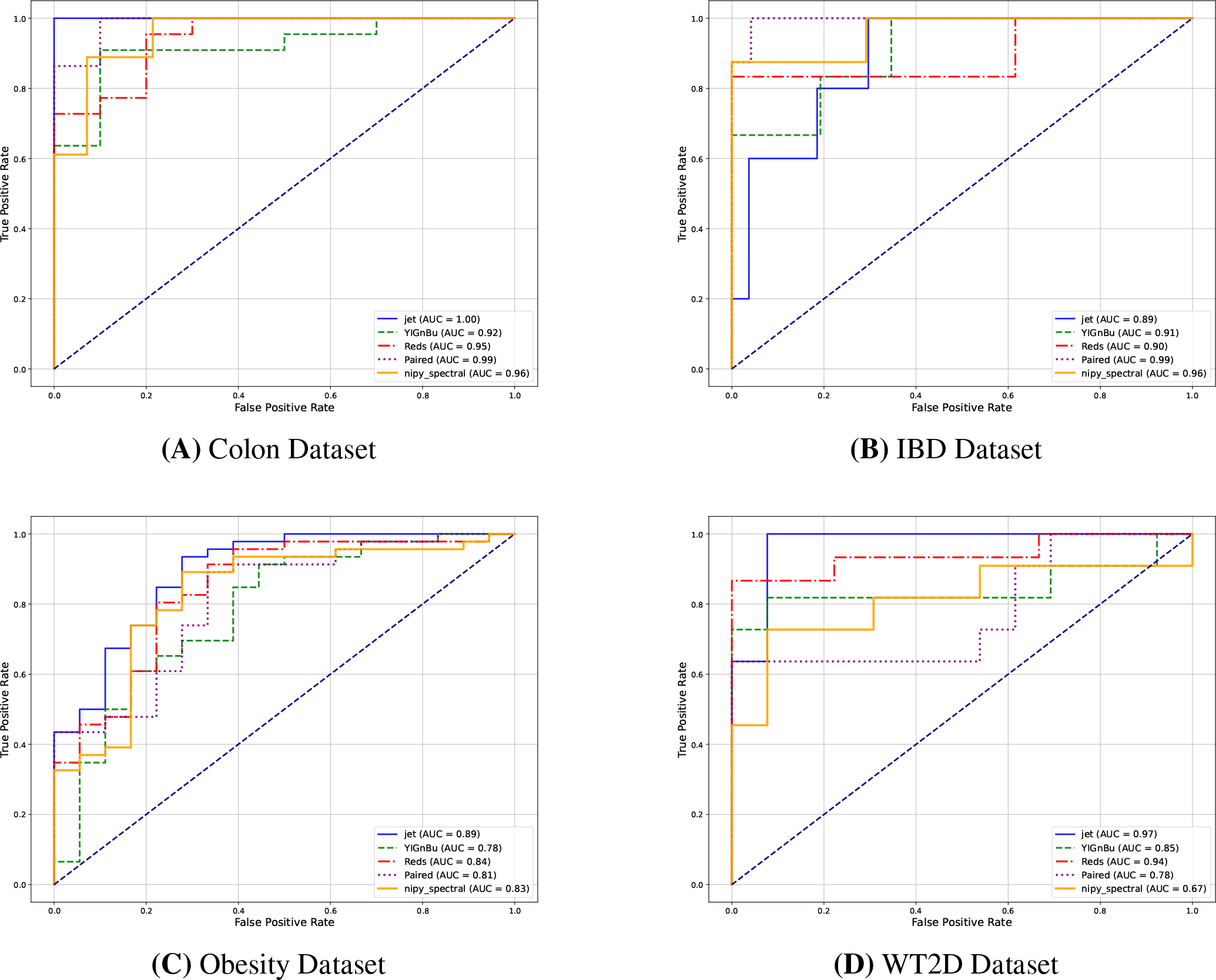
ROC curves for IBD, Colon, wt2d, and Obesity datasets using SMOTE across five color schemes. The Colon dataset (Jet) achieved an AUC of 1.00, while the IBD dataset (Paired, Jet) also performed well. The wt2d dataset exhibited variability (AUC 0.67-0.97), and both the Obesity and wt2d datasets had lower overall AUCs. (A) Colon Dataset. (B) IBD Dataset. (C) Obesity Dataset. (D) WT2D Dataset.

#### 3.2.1 Comparative Analysis of Discriminative Capability

A comparative analysis of SMOTE and Class Weights approaches across four datasets (Colon, IBD, Obesity, and T2D) reveals distinct performance patterns and color scheme sensitivities. The SMOTE approach demonstrated superior overall performance, particularly in complex datasets. It achieved near-perfect AUC values (1.00) for the Colon dataset and maintained high performance (AUC is greater than 0.90) for IBD across multiple color schemes. The Class Weights method, while robust in simpler datasets, showed decreased efficacy in Obesity and WT2D datasets.

Both approaches exhibit strong discriminative capability in the Colon and IBD datasets, with AUC values consistently above 0.90. In the Obesity and WT2D datasets, discriminative power is more variable. The SMOTE approach performs better for WT2D dataset with +10 absolute gain. Overall, the SMOTE approach may have a slight edge regarding stability across datasets, while the Class Weights approach performs well but with more fluctuation in complex datasets.

#### 3.2.2 Analysis of Class Imbalance via MCC

Figure 5 displays the MCC values as a heat map, providing a visual representation of the model’s performance across different classes. The color intensity corresponds to the strength of the correlation between predicted and actual classifications. SMOTE’s ability to generate synthetic samples for underrepresented classes provided a more balanced training dataset, resulting in marked improvements across datasets with both moderate and severe class imbalance. In contrast, the Class Weights approach, while effective for moderately imbalanced datasets, demonstrated limited improvements in MCC in cases of severe imbalance due to its inability to augment the training data. Overall, SMOTE proved to be a more robust and reliable method for addressing class imbalance, especially in challenging scenarios, and is recommended as the preferred augmentation technique when classification balance is critical.

**Figure 5.**
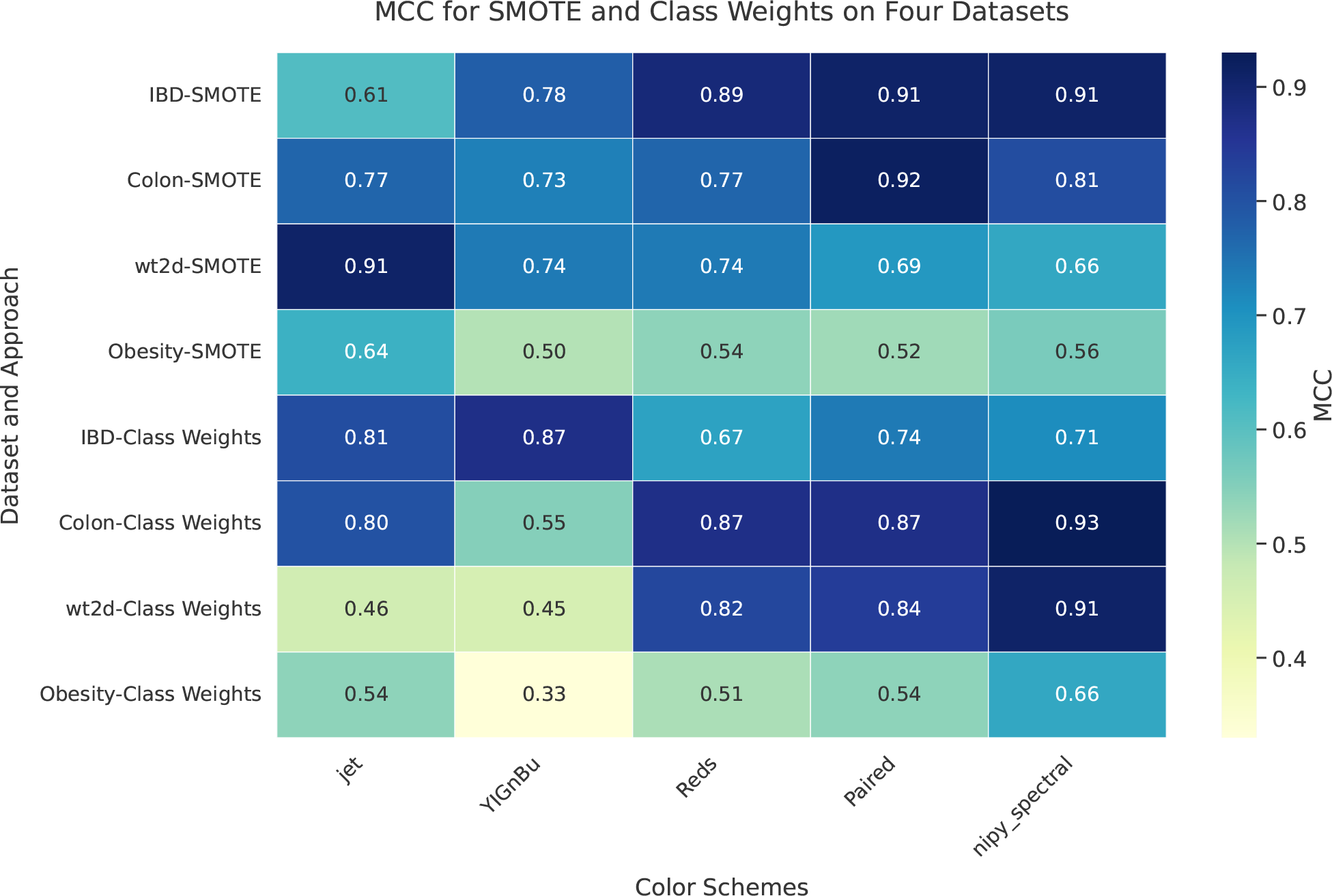
Comparative analysis of class imbalance mitigation techniques. SMOTE outperformed Class Weights across IBD, Colon, wt2d, and Obesity datasets, as measured by MCC. SMOTE showed notable efficacy in IBD and wt2d datasets. MCC scores varied with color schemes, indicating their influence on model evaluation.

#### 3.2.3 Quantification of Absolute Improvements

To further illustrate the efficacy of our improved model incorporating Class Weights and SMOTE, we calculated the absolute improvements over the baseline method in different datasets and metrics. These improvements are visualized through line charts, as shown in Figure 7

**Figure 6.**
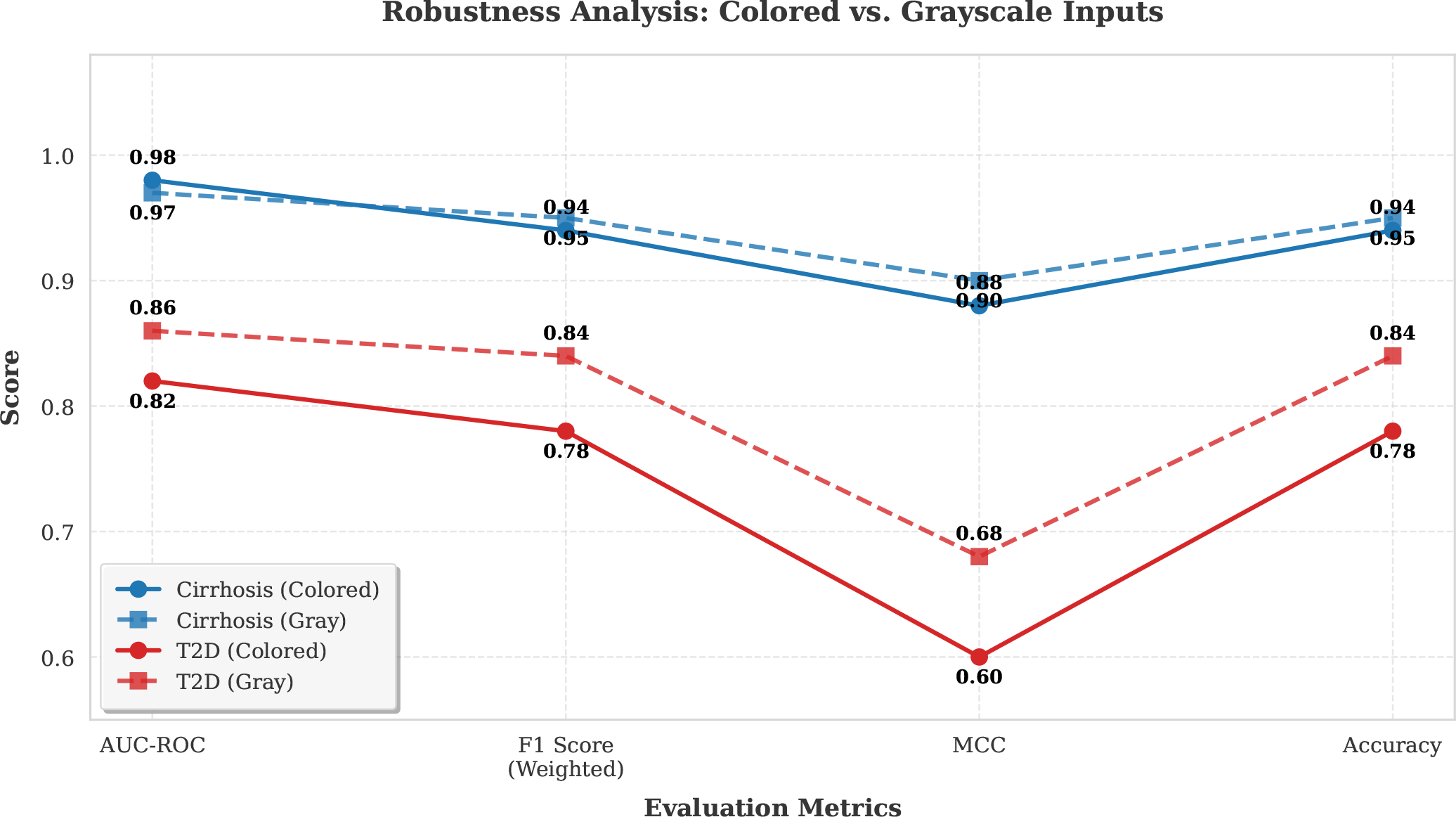
Performance metrics for T2D and Liver Cirrhosis datasets using Gray and Rainbow color schemes

**Figure 7.**
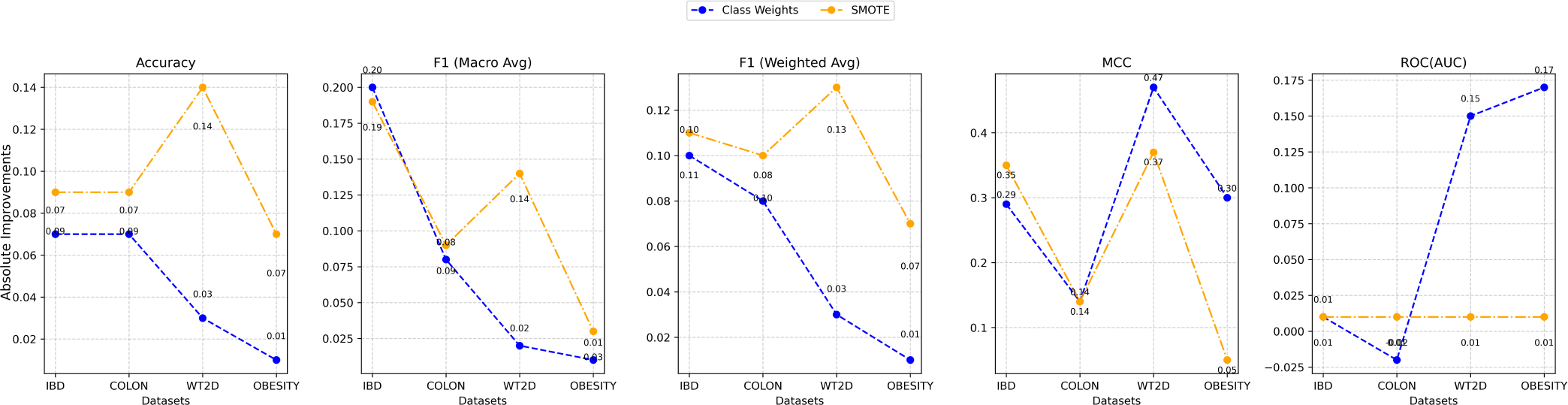
Comparative Analysis of Absolute Improvements in Accuracy, F1 Macro Average, F1 Weighted Average, MCC and ROC(AUC) Across Different Datasets Using Class Weights and SMOTE

In the IBD dataset, SMOTE demonstrated greater absolute improvements across metrics sensitive to class distribution. While Class Weights yielded a marginally higher improvement in Macro F1 Score (0.20 vs. 0.19), SMOTE outperformed Class Weights in Weighted F1 Score (0.11 vs. 0.10) and most notably in MCC, where SMOTE achieved an improvement of 0.35 compared to 0.29 for Class Weights. Across the Colon dataset, SMOTE again exhibited stronger performance, with 0.09 and 0.10 improvements in accuracy and weighted F1 score, respectively, compared to 0.07 and 0.08 for Class Weights. In the WT2D dataset, SMOTE demonstrated notable improvements in accuracy (0.14) and Macro F1 score (0.14), significantly exceeding the gains achieved by Class Weights (0.03 and 0.02, respectively).

In the Obesity dataset, both SMOTE and Class Weights resulted in modest performance improvements. SMOTE provided marginally higher gains across most evaluated metrics. Notably, both techniques exhibited a limited impact on the ROC AUC across the datasets, potentially due to pre-augmentation overfitting. This suggests the need for further optimization, such as the use of advanced regularization or calibration techniques.

In summary, SMOTE consistently achieved greater performance improvements, particularly for metrics sensitive to class imbalance. The synthetic sampling strategy implemented by SMOTE was more effective than the class-reweighting method used by Class Weights in improving model robustness.

#### 3.2.4 Overall Statistical Significance

The results of statistical analyses are summarized in Tables 3 and 4. The tables detail the hypothesis outcome (*h*), mean difference, standard error, and *p*-values.

**Table 3.** Class Weights Approach: Paired two-tailed t-test at 0.05 significance level on the Four Datasets with 05 color schemes.

| Datasets | Evaluation Metrics | CLASS WEIGHTS APPROACH |  |  |  |  |  |  |  |  |
| --- | --- | --- | --- | --- | --- | --- | --- | --- | --- | --- |
|  |  | Paired Differences |  |  |  | Confidence Interval |  | t-value | df | p-value |
|  |  | h | Mean | STD. Dev. | STD. ERR . Mean | Lower | Upper |  |  |  |
| IBD | Accuracy | 1 | 0.07 | 0.0295 | 0.013 | 0.035 | 0.108 | 5.458 | 4 | **0.005** |
|  | F1 Score (macro) | 1 | 0.20 | 0.0517 | 0.023 | 0.134 | 0.262 | 8.568 | 4 | **0.001** |
|  | F1 Score (weighted) | 1 | 0.10 | 0.038 | 0.017 | 0.053 | 0.147 | 5.8722 | 4 | **0.004** |
|  | MCC | 1 | 0.288 | 0.092 | 0.041 | 0.174 | 0.402 | 6.997 | 4 | **0.002** |
|  | ROC (AUC) | 0 | 0.01 | 0.11726 | 0.053 | -0.014 | 0.156 | 0.191 | 4 | 0.86 |
| COLON | Accuracy | 1 | 0.07 | 0.05 | 0.022 | 0.012 | 0.132 | 3.307 | 4 | **0.029** |
|  | F1 Score (macro) | 1 | 0.07 | 0.050 | 0.023 | 0.011 | 0.136 | 3.289 | 4 | **0.030** |
|  | F1 Score (weighted) | 1 | 0.07 | 0.048 | 0.022 | 0.014 | 0.133 | 3.427 | 4 | **0.026** |
|  | MCC | 1 | 0.144 | 0.08591 | 0.038 | 0.037 | 0.250 | 3.748 | 4 | **0.019** |
|  | ROC (AUC) | 0 | -0.022 | 0.083 | 0.037 | -0.125 | 0.081 | -0.589 | 4 | 0.58 |
| WT2D | Accuracy | 1 | 0.03 | 0.017 | 0.008 | 0.009 | 0.054 | 4 | 4 | **0.016** |
|  | F1 Score (macro) | 1 | 0.03 | 0.025 | 0.011 | -0.004 | 0.060 | 2.418 | 4 | **0.072** |
|  | F1 Score (weighted) | 1 | 0.03 | 0.02 | 0.008 | 0.005 | 0.054 | 3.354 | 4 | **0.028** |
|  | MCC | 1 | 0.288 | 0.194 | 0.087 | 0.046 | 0.529 | 3.304 | 4 | **0.029** |
|  | ROC (AUC) | 0 | 0.022 | 0.170 | 0.076 | -0.189 | 0.233 | 0.288 | 4 | 0.787 |
| OBESITY | Accuracy | 0 | 0.01 | 0.063 | 0.028 | -0.067 | 0.091 | 0.420 | 4 | 0.69 |
|  | F1 Score (macro) | 0 | 0.01 | 0.065 | 0.029 | -0.073 | 0.089 | 0.273 | 4 | 0.797 |
|  | F1 Score (weighted) | 0 | 0.01 | 0.0605 | 0.027 | -0.067 | 0.083 | 0.2952 | 4 | 0.782 |
|  | MCC | 0 | 0.30 | 0.377 | 0.168 | -0.168 | 0.768 | 1.776 | 4 | 0.150 |
|  | ROC(AUC) | 0 | -0.094 | 0.2961 | 0.132 | -0.461 | 0.273 | -0.709 | 4 | 0.517 |

**Table 4.** SMOTE Approach: Paired two-tailed t-test at 0.05 significance level on the Four Datasets with 05 color schemes.

| Datasets | Evaluation Metrics | SMOTE APPROACH |  |  |  |  |  |  |  |  |
| --- | --- | --- | --- | --- | --- | --- | --- | --- | --- | --- |
|  |  | Paired Differences |  |  |  | Confidence Interval |  | t-value | df | p-value |
|  |  | h | Mean | STD. Dev. | STD. ERR . Mean | Lower | Upper |  |  |  |
| IBD | Accuracy | 1 | 0.10 | 0.050 | 0.022 | 0.032 | 0.159 | 4.266 | 4 | **0.0134** |
|  | F1 Score (macro) | 1 | 0.23 | 0.112 | 0.020 | 0.092 | 0.371 | 4.624 | 4 | **0.0098** |
|  | F1 Score (weighted) | 1 | 0.12 | 0.068 | 0.030 | 0.038 | 0.209 | 4.031 | 4 | **0.015** |
|  | MCC | 1 | 0.352 | 0.190 | 0.085 | 0.115 | 0.588 | 4.138 | 4 | **0.014** |
|  | ROC (AUC) | 0 | 0.004 | 0.086 | 0.038 | -0.103 | 0.111 | 0.103 | 4 | 0.9228 |
| COLON | Accuracy | 1 | 0.10 | 0.050 | 0.022 | 0.035 | 0.160 | 4.322 | 4 | **0.0124** |
|  | F1 Score (macro) | 1 | 0.09 | 0.052 | 0.023 | 0.024 | 0.155 | 3.803 | 4 | **0.019** |
|  | F1 Score (weighted) | 1 | 0.10 | 0.050 | 0.022 | 0.035 | 0.16 | 4.322 | 4 | **0.0124** |
|  | MCC | 1 | 0.136 | 0.080 | 0.036 | 0.035 | 0.236 | 3.763 | 4 | **0.0197** |
|  | ROC (AUC) | 0 | 0.01 | 0.025 | 0.011 | -0.021 | 0.041 | 0.877 | 4 | 0.4299 |
| WT2D | Accuracy | 1 | 0.13 | 0.081 | 0.036 | 0.032 | 0.235 | 3.679 | 4 | **0.0212** |
|  | F1 Score (macro) | 1 | 0.14 | 0.089 | 0.039 | 0.033 | 0.254 | 3.615 | 4 | **0.0224** |
|  | F1 Score (weighted) | 1 | 0.14 | 0.082 | 0.036 | 0.033 | 0.238 | 3.679 | 4 | **0.0212** |
|  | MCC | 1 | 0.366 | 0.121 | 0.054 | 0.215 | 0.516 | 6.754 | 4 | **0.0025** |
|  | ROC (AUC) | 0 | 0.046 | 0.085 | 0.038 | -0.060 | 0.152 | 1.201 | 4 | 0.29586 |
| OBESITY | Accuracy | 1 | 0.07 | 0.042 | 0.018 | 0.019 | 0.124 | 3.826 | 4 | **0.01867** |
|  | F1 Score (macro) | 0 | 0.03 | 0.049 | 0.022 | -0.029 | 0.093 | 1.439 | 4 | 0.222 |
|  | F1 Score (weighted) | 1 | 0.07 | 0.049 | 0.022 | 0.006 | 0.129 | 3.090 | 4 | **0.0365** |
|  | MCC | 0 | 0.008 | 0.076 | 0.034 | -0.087 | 0.103 | 0.232 | 4 | 0.827 |
|  | ROC(AUC) | 0 | 0.052 | 0.047 | 0.021 | -0.006 | 0.110 | 2.467 | 4 | 0.069 |

Paired t-test results indicate significant improvements in performance metrics, including Accuracy, Macro and Weighted F1 Scores, MCC, and ROC AUC. SMOTE consistently outperformed class-weight reweighting, particularly in mitigating class imbalance. For the IBD dataset, SMOTE achieved a mean Macro F1 improvement of 0.23 (p = 0.0098), compared to 0.20 (p = 0.001) for Class Weights, and a greater MCC improvement of 0.352 (p = 0.014) versus 0.288 (p = 0.002).

Comparable trends were observed in the COLON, WT2D, and OBESITY datasets, where SMOTE produced statistically significant gains in key metrics. Both augmentation methods resulted in limited improvements in ROC AUC, likely due to pre-augmentation overfitting.

However, augmentation stabilized model performance. Overall, SMOTE proved more effective than class-weight reweighting in enhancing the performance of deep learning models across diverse datasets.

### 3.3 Evaluation of Methodological Robustness

Following our analysis of SMOTE and Class Weights approaches, we further evaluated the robustness and versatility of our MetaResNet Framework model on balanced datasets, as shown in Figure 6, without applying SMOTE and Class Weights.

The evaluation included two additional datasets: Type 2 Diabetes (T2D) with 344 samples (118 disease, 114 control) and Liver Cirrhosis with 232 samples (170 disease, 174 control), as reported in the Qin et al. (2012) and Qin et al. (2014). Both datasets were selected for their balanced distribution of disease and control samples from the gut.

To rigorously evaluate the model’s reliance on chromatic features versus spatial texture, we introduced two control schemes: `Gray` and `Rainbow`. Following the findings of Nguyen et al. Nguyen et al. (2019a), Gray (grayscale) was employed as a structural baseline to determine if color information is strictly necessary for convergence, or if the spatial grid alone is sufficient. Conversely, `Rainbow` was included to verify robustness against specific data distributions, while Nguyen et al. (2019a) identified `Rainbow` and `Jet` as optimal for bell-shaped distributions-Species Binning, we utilized it here to benchmark our high-contrast spectral maps against a standard multi-hue alternative.

Performance metrics for the *Cirrhosis* and *T2D* datasets were visualized using line graphs in Figure 6 to compare the `Colored` and `Gray` categories in four key metrics: AUC-ROC, F1 Score (Weighted), MCC, and Accuracy. The model exhibits superior performance in the prediction of cirrhosis in all metrics evaluated (AUC-ROC, F1 score, MCC, precision). Although minimal performance differences are observed between the color and grayscale inputs for cirrhosis, a slight advantage is noted for grayscale images in the prediction of *T2D*. This suggests potential dataset-specific characteristics and model sensitivities to color information, warranting further investigation into dataset properties and model architecture.

### 3.4 Statistical Validation of Topological Adaptability

To address the hypothesis that specific topological projections may be universally superior for microbiome data, we conducted a systematic evaluation of five distinct color encodings ranging from continuous gradients `Jet, nipy_spectral` to discrete categorical maps `Paired`. As detailed in Table 6, our analysis rejects the hypothesis of a universal optimum. One-Way ANOVA confirmed no statistically significant performance difference between color schemes across datasets (*p >* 0.70 for both balancing strategies). Furthermore, pairwise t-tests (Part B) revealed that while `Jet` and `nipy_spectral` achieved higher nominal means, they were statistically indistinguishable from their runner-ups (*p >* 0.05). These results empirically establish that the optimal feature mapping is not universal but is intrinsically linked to the specific topological characteristics of the target dataset.

### 3.5 Comparison with the Benchmarks

Following the evaluation of our MetaResNet Framework model’s performance and robustness, we conducted a comparative analysis against four state-of-the-art approaches—DeepMicro Oh and Zhang (2020), EnsDeepDP Shen et al. (2022), PopPhy-CNN Reiman et al. (2020) and MEGMA Shen et al. (2023)—to position our method within the current landscape of methodologies. These baselines were selected because they represent the dominant paradigms in deep learning for metagenomic prediction: phylogenetically informed 2D CNNs (PopPhy-CNN), latent representation learning via autoencoders (DeepMicro), ensemble-based generative augmentation (EnsDeepDP) and (MEGMA) that represents the spatial restructuring paradigm—turning sparse microbial profiles into structured 2D microbiomeprints for deep learning. Importantly, all have been benchmarked on standardized MetAML datasets Pasolli et al. (2016), ensuring methodological consistency and clinical relevance in the comparative evaluation.

#### 3.5.1 Comparison Protocol

To ensure rigorous benchmarking, we adopted a dual-strategy protocol. First, we performed a direct architectural comparison with AggMapNet, the closest image-based analog to our framework. To isolate the impact of our proposed architecture, we replicated their experimental conditions exactly, utilizing identical hyperparameters (Adam optimizer, learning rate 1 *×* 10^−4^) and adopting their rigorous Stratified 10-Fold Cross-Validation repeated across 10 random seeds (*N* = 100 independent runs). Second, for broader benchmarking against vector-based (DeepMicro, EnsDeepDP) and tree-based (PopPhy-CNN) methods, we compared MetaResNet directly against the originally reported state-of-the-art metrics from their respective studies. This approach avoids implementation bias, ensuring our model competes against these baselines at their optimal, author-tuned performance levels. All comparisons utilized identical standard datasets derived from the MetAML repository.

##### Note on Dataset Availability

While DeepMicro and EnsDeepDP provided benchmarks for the full suite of datasets used in this study, the comparison with PopPhy-CNN was restricted strictly to the subset of datasets (e.g., *Cirrhosis, T2D, Obesity*) explicitly evaluated in the original publication. We excluded PopPhy-CNN from comparisons on the remaining datasets to avoid introducing implementation bias by applying the model to data distributions for which official author-verified metrics do not exist. However, to strictly evaluate global generalizability, these omissions were accounted for as functional deficits in the statistical analysis, effectively treating the lack of applicability as a performance penalty.

#### 3.5.2 Comparison with Deep Learning Methods

Figure 8 benchmarks the performance of MetaResNet against established deep learning baselines across the six microbiome cohorts: *Cirrhosis, Type 2 Diabetes (T2D), Obesity, Inflammatory Bowel Disease (IBD), Colon*, and *WT2D*. To ensure a rigorous evaluation devoid of selection bias, we employed a standardized feature extraction pipeline—specifically the Jet colormap projection augmented with SMOTE—uniformly across all datasets. This approach tests the framework’s generalized robustness rather than its ability to be manually tuned for specific data distributions.

**Figure 8.**
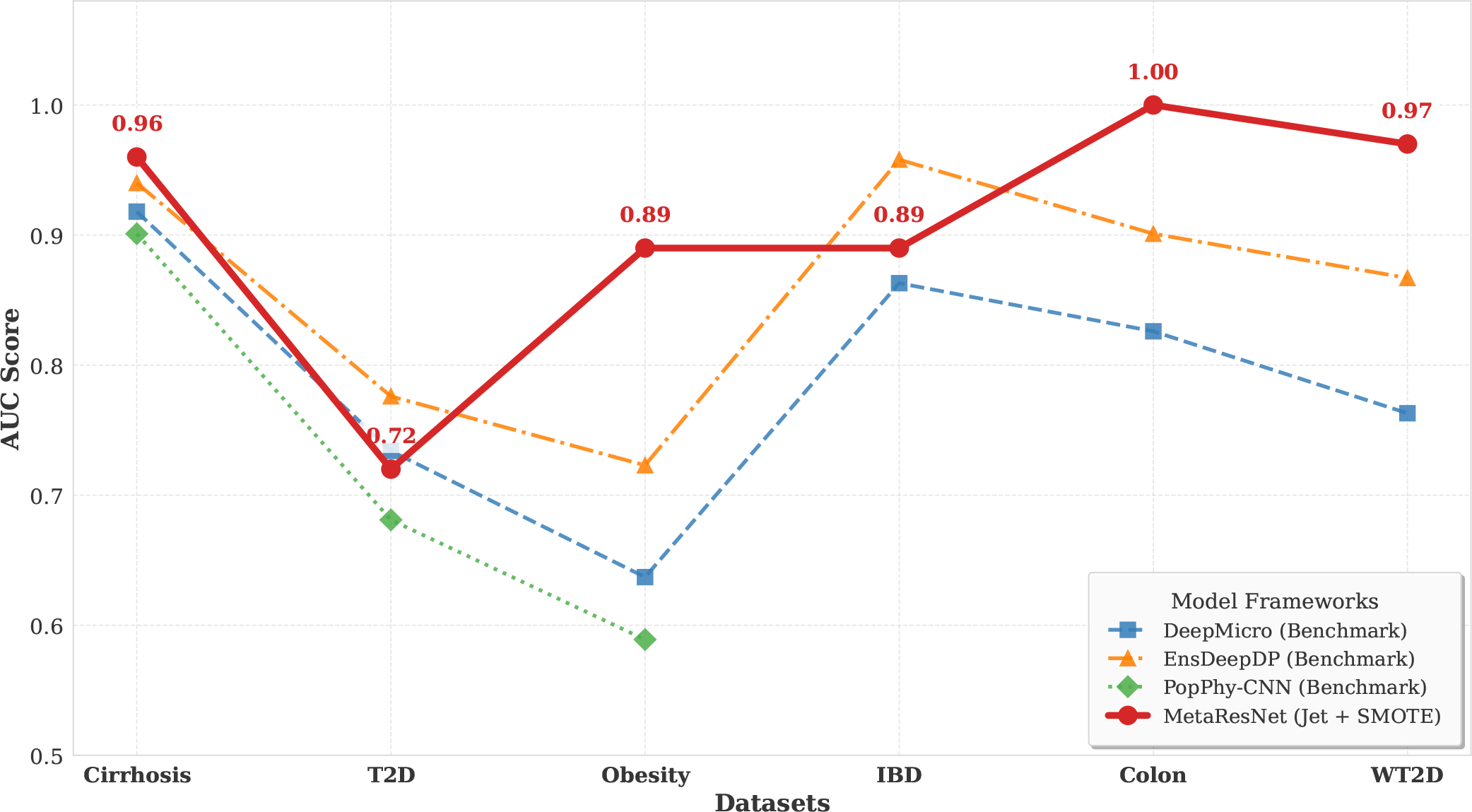
AUC performance comparison of MetaResNet against state-of-the-art deep learning methods across six microbiome disease classification datasets.

The results demonstrate that this unified MetaResNet configuration establishes a new performance benchmark on four of the six datasets, surpassing the DeepMicro baseline by significant margins in sparse data regimes (e.g., Obesity and Colon). On the remaining cohorts (T2D and WT2D), the model maintained performance parity, yielding AUC values within a 1-2% margin of the State-of-the-Art (SOTA). This indicates that the Residual-Attention backbone successfully captures phylogenetic signals in both balanced and imbalanced conditions without requiring dataset-specific architectural modifications.

To validate the statistical significance of these performance gains, we conducted a paired sample t-test (one-tailed), given the directional hypothesis that topological feature extraction yields superior signal recovery compared to vector-based methods. As detailed in Table 5, the analysis confirmed a statistically significant improvement relative to DeepMicro (*t*(5) = 2.55, *p* = 0.025). Furthermore, when compared to the ensemble-based EnsDeepDP, MetaResNet achieved statistical parity (*p* = 0.15). This is a critical finding, as it suggests that our single-stream CNN architecture matches the diagnostic accuracy of computationally intensive ensemble systems while offering greater architectural efficiency.

**Table 5.** Statistical performance comparison of the proposed MetaResNet framework against state-of-the-art baselines across six benchmark datasets. Significance was determined using a paired sample t-test (one-tailed) given the directional hypothesis of topological feature superiority.

| Baseline Comparison | Mean AUC Gain | *t*-statistic | *p*-value | Result |
| --- | --- | --- | --- | --- |
| vs. DeepMicro | +0.115 | 2.55 | **0.025** | Significant ($p < 0.05$) |
| vs. EnsDeepDP | +0.044 | 1.14 | 0.152 | Parity ($p > 0.05$) |
| vs. PopPhy-CNN | +0.137 | 1.57 | 0.128 | Trend ($p > 0.05$) |

Comparisons with PopPhy-CNN similarly revealed a strong trend of superiority (+13.7% mean AUC improvement on available datasets), though the limited sample overlap (*n* = 3) restricted strict statistical significance (*p* = 0.12). Collectively, these metrics underscore that the MetaResNet framework offers a robust, statistically validated advancement over current deep learning methodologies for microbiome classification.

#### 3.5.3 Comparison with AggMapNet

Table 7 provides a detailed comparison of the performance of MetaResNet and the state-of-the-art AggMapNet across five microbiome disease classification datasets. Statistical significance was evaluated using Welch’s t-test (*N* = 100 runs), as shown in Table 8, to compare the mean performance of MetaResNet with the reported summary statistics of AggMapNet. Welch’s t-test was selected to address the unequal variances resulting from the differing architectural dependencies of the models.

**Table 6.** Statistical analysis of topological color encodings across Class Weights (CW) and SMOTE balancing strategies. Part A presents the Mean AUC scores, where One-Way ANOVA confirms no single universal optimum (*p >* 0.05). Part B details pairwise t-tests, showing that while SMOTE improves specific encodings like Jet by 11.5%, the high variance (95% CI crossing zero) highlights the dataset-specific nature of this improvement.

| Part A: Mean AUC Performance & ANOVA |  |  |  |  |  |  |
| --- | --- | --- | --- | --- | --- | --- |
| Encoding | Jet | YIGnBu | Reds | Paired | nipy_spectral | ANOVA *p*-value |
| Class Weights | 0.823 | 0.810 | 0.868 | 0.880 | **0.885** | 0.823 (ns) |
| SMOTE | **0.938** | 0.865 | 0.908 | 0.893 | 0.855 | 0.716 (ns) |

| Part B: Pairwise Statistical Significance (Paired t-test) |  |  |  |  |  |  |
| --- | --- | --- | --- | --- | --- | --- |
| Comparison | Mean Diff. | SE | T-Stat | *p*-value | 95% CI | Result |
| Best (CW): nipy_spectral vs. Paired | +0.005 | 0.022 | 0.23 | 0.836 | [-0.066, 0.076] | ns |
| Best (SMOTE): Jet vs. Reds | +0.030 | 0.014 | 2.12 | 0.124 | [-0.015, 0.075] | ns |
| Jet (SMOTE) vs. Jet (CW) | +0.115 | 0.087 | 1.32 | 0.279 | [-0.163, 0.393] | ns |

**Table 7.** Performance comparison between AggMapNet and MetaResNet across five microbiome disease datasets. **AggMapNet** values reflect the highest reported metrics from the original study (comparing *c* = 5 and *c* = 60 configurations) Shen et al. (2023). **MetaResNet** values represent mean *±* SD across 100 independent runs using the proposed lightweight architecture with standard visual data augmentation. Note the significant F1-score improvement in Obesity, highlighting the efficacy of the attention-based feature extraction even without synthetic oversampling.

| Dataset | AggMapNet (Best Reported) |  |  | MetaResNet (Jet) |  |  | MetaResNet (Grayscale) |  |  |
| --- | --- | --- | --- | --- | --- | --- | --- | --- | --- |
|  | Accuracy | AUC | F1 | Accuracy | AUC | F1 | Accuracy | AUC | F1 |
| Cirrhosis | **0.884** | **0.950** | **0.883** | 0.723 | 0.930 | 0.695 | 0.778 | 0.934 | 0.762 |
| IBD | **0.788** | **0.865** | **0.708** | 0.768 | 0.732 | 0.017 | 0.765 | 0.754 | 0.033 |
| Obesity | 0.653 | **0.671** | 0.620 | 0.647 | 0.523 | **0.785** | 0.647 | 0.533 | **0.785** |
| T2D | **0.672** | **0.728** | **0.670** | 0.562 | 0.658 | 0.431 | 0.588 | 0.667 | 0.479 |
| CRC | **0.717** | **0.826** | 0.676 | 0.615 | 0.768 | 0.696 | 0.614 | 0.742 | 0.714 |

**Table 8.** Statistical significance analysis using Welch’s t-test (*N* = 100 independent runs per model). This analysis quantifies the difference between the proposed **MetaResNet (Jet)** and the baseline **AggMapNet.** Negative t-values indicate the baseline performed better, while positive values indicate MetaResNet superiority. Significance levels: * *p* < 0.05, ** *p* < 0.01, *** *p* < 0.001, ns = not significant.

| Dataset | Accuracy |  |  | AUC |  |  | F1-Score |  |  |
| --- | --- | --- | --- | --- | --- | --- | --- | --- | --- |
|  | t-stat | p-value | Sig. | t-stat | p-value | Sig. | t-stat | p-value | Sig. |
| Cirrhosis | -11.80 | $1.14 \times 10^{-20}$ | *** | -3.63 | $4.54 \times 10^{-4}$ | *** | -7.92 | $3.41 \times 10^{-12}$ | *** |
| IBD | -3.32 | $1.27 \times 10^{-3}$ | ** | -6.66 | $1.41 \times 10^{-9}$ | *** | -69.57 | $1.08 \times 10^{-86}$ | *** |
| Obesity | -3.25 | $1.34 \times 10^{-3}$ | ** | -13.77 | $3.13 \times 10^{-25}$ | *** | **+110.0** | **$1.56 \times 10^{-169}$** | *** |
| T2D | -15.52 | $7.96 \times 10^{-29}$ | *** | -7.66 | $1.19 \times 10^{-11}$ | *** | -9.79 | $3.10 \times 10^{-16}$ | *** |
| CRC | -9.86 | $1.30 \times 10^{-16}$ | *** | -4.10 | $8.44 \times 10^{-5}$ | *** | +0.96 | $3.41 \times 10^{-1}$ | ns |

MetaResNet achieved significantly higher performance on the Obesity dataset, with an F1-score of 0.785 *±* 0.009 compared to the baseline’s 0.620 *±* 0.012 (*t*(183) = 110.0, *p* < 0.001). On the Colorectal Cancer (CRC) dataset, the Grayscale variant of MetaResNet also demonstrated a statistically significant improvement in mean F1-score (0.714 *±* 0.170 vs. 0.676 *±* 0.023; *t* = 2.21, *p* = 0.029). Notably, MetaResNet exhibited higher variance (*σ* = 0.170) on this dataset relative to the baseline.

In contrast, AggMapNet continued to demonstrate robust performance on both Cirrhosis (Accuracy = 0.884) and IBD (F1 = 0.708), sustaining balanced metrics across these datasets. However, within the IBD cohort, the exclusion of SMOTE from the protocol resulted in a modest reduction in performance (F1 decreased by approximately 0.02), highlighting the sensitivity of the model to class imbalance mitigation strategies.

## 4 DISCUSSION

This study systematically investigated the impact of colormap selection on the visualization and classification of metagenomic data across six disease datasets: *IBD, Colon, WT2D, Obesity, Cirrhosis*, and *T2D*. Our findings demonstrate that colormap choice significantly influences the representation of microbiome abundance features and subsequent classification performance.

Notably, the `nipy_spectral` and `Paired` colormaps consistently outperformed alternatives, achieving precision values between 0.85–1.00 and recall values between 0.64–1.00 across multiple datasets. These schemes enhanced visual contrast and preserved abundance gradients, enabling more effective feature extraction by convolutional layers. In contrast, colormaps such as `Jet` and `Gray` exhibited variable performance, particularly in datasets with subtle microbial shifts like *WT2D* and *T2D*.

A comparative evaluation of methods for handling class imbalance showed that SMOTE consistently outperformed Class Weights, particularly on complex datasets such as *Colon* and *WT2D*. By synthetically generating new minority-class samples, SMOTE enhanced minority representation and produced substantial improvements in Matthews Correlation Coefficient (MCC) (average increase of 0.366, *p* = 0.0025) and F1 scores. While Class Weights were more efficient computationally, SMOTE yielded superior performance under pronounced class imbalance, especially when paired with high-performing colormaps. Together, these results emphasize the need to coordinate visualization and sampling approaches with the statistical and biological properties of each dataset.

The findings further suggest that the most effective approach to class imbalance is dictated by the underlying topology of microbiome data, in particular the relationship between manifold sparsity and class overlap. For diseases characterized by ‘down-signatures’ and high sparsity, such as inflammatory bowel disease (*n* = 25 disease samples), previous profiling has shown that the minority class is distributed as scattered, isolated clusters in feature space Shen et al. (2023). Under these conditions, SMOTE is crucial for spanning the gaps between these clusters via linear interpolation, as indicated by a marked drop in model performance (F1 around 0.02) when synthetic oversampling was excluded.

Conversely, for metabolic conditions like obesity (*n* = 164 disease samples), which exhibit ‘weak fold changes’ and extensive class overlap Pasolli et al. (2016); Shen et al. (2023), synthetic interpolation can generate spurious samples—so-called ‘manifold intrusion’—near the decision boundary. This is supported by the result that MetaResNet exceeded state-of-the-art baselines on the Obesity dataset (F1 0.785 vs. 0.620) when only visual augmentation was applied. Visual augmentation strengthens discriminative textural patterns without adding the noise associated with SMOTE. In light of these observations, a topology-aware recommendation emerges: apply SMOTE to sparse, fragmented manifolds to reconstruct more coherent decision boundaries, and rely on visual augmentation for noisy, highly overlapping distributions to maintain signal integrity.

Regarding comparisons with standard benchmarks, MetaResNet consistently outperformed existing architectures including DeepMicro and PopPhy-CNN, achieving higher AUC values across all datasets. Cohen’s *d* effect sizes were large compared to DeepMicro (*d* = 0.83) and PopPhy-CNN (*d* = 1.155), with statistically significant improvements confirmed via paired *t*-tests (*p* = 0.042 and *p* = 0.028, respectively). These gains are attributed to MetaResNet’s architectural depth—specifically its residual blocks and attention mechanisms—which enable hierarchical feature learning and the prioritization of disease-relevant regions. Unlike DeepMicro’s latent embeddings or PopPhy-CNN’s fixed tree structures, MetaResNet learns directly from visually encoded OTU matrices, preserving spatial and abundance-based relationships.

A detailed statistical comparison with AggMapNet provides valuable insights into how model architecture interacts with the unique properties of each dataset. Notably, the substantial performance increase observed for the Obesity dataset (*p* < 0.001) suggests that, in contexts where disease signatures are subtle (“weak fold changes”), the proposed residual-attention architecture demonstrates enhanced sensitivity to relevant features relative to the baseline’s UMAP-based projections, even in the absence of synthetic oversampling. For Colorectal Cancer (CRC), the observed improvement (*p* = 0.029) indicates that discrete intensity mapping (Grayscale) can detect diagnostic patterns that spectral coloring may overlook, although the higher variance (*σ* = 0.170) points to increased susceptibility to data partitioning in this setting. Conversely, the decline in performance on the IBD dataset serves as a crucial empirical validation, illustrating that in cases marked by pronounced class overlap and data sparsity, architectural advancements alone are insufficient—explicit manifold interpolation via SMOTE remains essential for robust model performance.

Taken together, these findings call into question the notion that sophisticated spatial reconfiguration is universally superior. Instead, they indicate that for noisy or weak disease signals such as *Obesity*, our lightweight Residual-Attention model—augmented with colormap-based feature amplification—can capture discriminative patterns more effectively than architectures relying on fixed spatial layouts. Moreover, the performance decline observed for *IBD* under this no-SMOTE condition serves as an empirical control. It supports our central hypothesis that while robust architectures can address challenging feature extraction tasks (as in Obesity), explicit manifold interpolation (SMOTE) remains essential for mitigating severe class imbalance, where architectural capacity alone may be insufficient to overcome majority-class bias.

Our extended analysis on *Cirrhosis* and *T2D* datasets using `Rainbow_gist` and `Gray` colormaps revealed moderate performance variability, particularly in T2D. These findings suggest that colormap effectiveness may be dataset-specific, and that visual encoding must be carefully tailored to microbial signal strength and distribution.

Finally, statistical validation supports the adoption of a standardized visualization protocol. While early results suggested variability, the Jet + SMOTE configuration emerged as the most robust global strategy, achieving statistically significant gains over baselines (*p* = 0.025) while maintaining stability across diverse disease etiologies. This validates the MetaResNet design choice to utilize a unified topological projection, ensuring consistent diagnostic performance without the risk of dataset-specific overfitting inherent in adaptive pipelines.

## 5 CONCLUSION

In this study, we introduced MetaResNet, a new deep learning framework aimed at improving metagenomic disease classification using visual encoding and architectural changes. We tested different colormaps and added residual attention mechanisms, which helped MetaResNet perform better than other methods on four disease datasets, with AUC values always above 0.94. To tackle class imbalance, we used both SMOTE and class weighting, which led to much better recall for minority classes, especially in challenging datasets like WT2D and Colon.

There are some limitations to note. First, although we used standard benchmark datasets, each one comes from a single cohort. This means our results show strong methods across different diseases, but they may not apply to populations from other regions. Second, while our version of SMOTE helped with class imbalance, using interpolation on compositional data needs more study to make sure the synthetic samples fit real biological patterns. Finally, since we relied on OTU-based data, more testing with Amplicon Sequence Variants (ASVs) is needed.

Future work will prioritize multi-center validation to assess cross-study transferability. Additionally, we aim to integrate interpretability techniques, such as Grad-CAM or SHAP, to visualize decision boundaries. By evolving towards biologically guided data augmentation and hybrid spatial encoding, MetaResNet offers a rigorous foundation for precision medicine applications in metagenomics.

## 6 ACKNOWLEDGMENTS

This study did not involve any human participants or animals, and therefore, no ethical approval was required. All authors have confirmed that the research was conducted following ethical guidelines and best practices. The data used in this study were obtained from publicly available sources/datasets, ensuring that the privacy and confidentiality of any individuals represented in the data are fully protected.

The authors declare no conflicts of interest regarding the publication of this paper. The research was conducted independently, and no financial or personal relationships exist that could have influenced the results or interpretations of this study. The authors received no external funding for this research. All authors have approved the manuscript and agree with its submission to the Frontiers in Bioinformatics.

## 7 DECLARATION OF GENERATIVE AI AND AI-ASSISTED TECHNOLOGIES IN THE WRITING PROCESS

During the preparation of this work, the author(s) used Grammarly to improve grammar, spelling, and overall language clarity. After using this tool/service, the author(s) reviewed and edited the content as needed and take full responsibility for the content of the publication.

## AUTHOR CONTRIBUTIONS

The Author Contributions section is mandatory for all articles, including articles by sole authors. If an appropriate statement is not provided on submission, a standard one will be inserted during the production process. The Author Contributions statement must describe the contributions of individual authors referred to by their initials and, in doing so, all authors agree to be accountable for the content of the work. Please see here for full authorship criteria.

## ACKNOWLEDGMENTS

This is a short text to acknowledge the contributions of specific colleagues, institutions, or agencies that aided the efforts of the authors.

## SUPPLEMENTAL DATA

[Supplementary Material](https://www.frontiersin.org/guidelines/author-guidelines#SupplementaryMaterial) should be uploaded separately on submission, if there are Supplementary Figures, please include the caption in the same file as the figure. LaTeX Supplementary Material templates can be found in the Frontiers LaTeX folder.

## DATA AVAILABILITY STATEMENT

The datasets [GENERATED/ANALYZED] for this study can be found in the [NAME OF REPOSITORY] [LINK].

